# Distinct C_4_ Sub-Types and C_3_ Bundle Sheath Isolation In The Paniceae Grasses

**DOI:** 10.1101/162644

**Authors:** Jacob D. Washburn, Josh Strable, Patrick Dickinson, Satya S. Kothapalli, Julia M. Brose, Sarah Covshoff, Gavin C. Conant, Julian M. Hibberd, J. Chris Pires

## Abstract

In C_4_ plants, the enzymatic machinery underpinning photosynthesis can vary, with, for example, three distinct C_4_ acid decarboxylases being used to release CO_2_ in the vicinity of RuBisCO. For decades, these decarboxylases have been used to classify C_4_ species into three biochemical sub-types. However, more recently the notion that C_4_ species mix and match C_4_ acid decarboxylases has increased in popularity and, as a consequence, the validity of specific biochemical sub-types has been questioned. Using five species from the grass tribe Paniceae, we show that, while in some species transcripts encoding multiple C_4_ acid decarboxylases accumulate, in others, transcript abundance and enzyme activity is almost entirely from one decarboxylase. In addition, the development of a bundle sheath isolation procedure for a close C_3_ species in the Paniceae enables the preliminary exploration of C_4_ sub-type evolution.

## Introduction

C_4_ photosynthesis is often considered the most productive mechanism by which plants convert sunlight into chemical energy (Sage, 2004; Wang et al., 2012; Niklaus and Kelly, 2019; Kopriva and Weber, 2021). The C_4_ pathway leads to increased photosynthetic efficiency because high concentrations of CO_2_ are supplied to RuBisCO. Since its discovery in the 1960s (Hatch and Slack, 1966), a unified understanding of the biochemistry underpinning C_4_ photosynthesis has emerged. This basic system comprises a biochemical pump that initially fixes HCO_3_^−^ into C_4_ acids in mesophyll (M) cells. Subsequently, diffusion of these C_4_ acids into a separate compartment, followed by their decarboxylation, generates high concentrations of CO_2_ around RuBisCO. In many plants, the release of CO_2_ occurs in bundle sheath (BS) cells (Hatch, 1992; Furbank, 2016; von Caemmerer et al., 2017). Although this pump demands additional ATP inputs, in warm environments where RuBisCO catalyzes high rates of oxygenation (and therefore photorespiration), the C_4_ pathway increases photosynthetic efficiency compared with the ancestral C_3_ state.

Elucidation of the C_4_ pathway was initially based on analysis of sugarcane (*Saccharum spp.* L.) and maize (corn, *Zea mays* L.), which both use the chloroplastic NADP-DEPENDENT MALIC ENZYME (NADP-ME) to release CO_2_ in BS cells. However, it became apparent that not all species used this chloroplastic enzyme. For example, *Megathyrsus maximus* (formerly *Panicum maximum*)*, Urochloa texanum* (formerly *Panicum texanum*), and *Sporobolus poiretti* used the cytosolic enzyme PHOSPHONENOLPYRUVATE CARBOXYKINASE (PEPCK) (Edwards et al., 1971) to release CO_2_ in the BS, whereas *Atriplex spongiosa* and *Panicum miliaceum* showed high activities of the mitochondrial NAD-DEPENDENT MALIC ENZYME (NAD-ME) (Hatch and Kagawa, 1974). These findings led to the consensus that different C_4_ species made preferential use of one C_4_ acid decarboxylase and resulted in the classification of C_4_ plants into one of three distinct biochemical pathways (Edwards et al., 1971; Hatch et al., 1975; Hatch and Kagawa, 1976). According to Furbank (2016), there was some early discussion about whether the sub-types were mutually exclusive or if one species might employ two or more sub-types together, but in general, the sub-types were described as distinct (Hatch, 1987).

For several decades this description of three sub-types has been standard practice (Sheen, 1999; Hibberd and Covshoff, 2010) and even used in taxonomic classification (Brown, 1977). However, more recent work has provided evidence that some C_4_ species use multiple C_4_ acid decarboxylases. Maize, for example, was traditionally classified as using NADP-ME but evidence has mounted that it and sugar-cane both have high activities of PEPCK (Walker et al., 1997; Wingler et al., 1999; Majeran et al., 2010; Furbank, 2011; Pick et al., 2011; Bellasio and Griffiths, 2013; Sharwood et al., 2014; Wang et al., 2014; Koteyeva et al., 2015; Weissmann et al., 2016; Cacefo et al., 2019). This blurring of the NADP-ME C_4_ sub-type coincided with observations that many plants with high amounts of PEPCK also contained either NADP-ME of NAD-ME (Furbank, 2011). Furthermore, computational models of the C_4_ pathways suggested that BS energy requirements could not be met in a system with only PEPCK decarboxylation (Wang et al., 2014). It has therefore been suggested that PEPCK may never function on its own as a distinct sub-type (Furbank, 2011; Bräutigam et al., 2014; Wang et al., 2014).

Alternatives to the three sub-type classification have since been proposed and used in a number of recent publications. These include a two sub-type system (based on the use of NADP-ME or NAD-ME), as well as a four sub-type classification placing species into NADP-ME, NAD-ME, NADP-ME + PEPCK, and NAD-ME + PEPCK sub-types (Wang et al., 2014; Washburn et al., 2015; Rao and Dixon, 2016). At present, none of these classification schemes has been widely adopted by the community. Moreover, convincing experimental evidence (i.e., transcriptomic, or proteomic data) that species traditionally defined as belonging to the PEPCK sub-type actually use another C_4_ acid decarboxylation enzyme at a higher level than PEPCK is lacking, while enzyme activity measurements in the older literature indicate strong PEPCK predominance for several species (Gutierrez et al., 1974; Prendergast et al., 1987; Lin et al., 1993).

Only one group of species, the tribe Paniceae (Poaceae) has been documented to contain all three classical biochemical sub-types of C_4_ photosynthesis together in a pattern consistent with a single C_4_ origin (Sage et al., 2011). The subtribe Cenchrinae consists of species using the classical NADP-ME C_4_ sub-type, the subtribe Melinidinae the PEPCK sub-type, and the Panicinae the NAD-ME sub-type (Gutierrez et al., 1974; Prendergast et al., 1987; Lin et al., 1993). The sub-tribes Cenchrinae, Melinidinae, and Panicineae (CMP) form a well-supported phylogenetic clade of C_4_ species with many C_3_ species sister to the clade (Vicentini et al., 2008; Grass Phylogeny Working Group II, 2012; Washburn et al., 2015). Studies based solely or predominantly on nuclear genes have confirmed this CMP clade, but also placed the sub-tribe Anthephorineae as sister to the CMPA clade. These phylogenies would create a CMPA clade of C_4_ species potentially sharing a single C_3_ ancestor (Vicentini et al., 2008; Washburn et al., 2017). This clade is here referred to as the CMP(A) clade in order to indicate the incongruence found between nuclear and chloroplast phylogenies (Figure 1). The analyses here performed would be equally valid regardless on the inclusion of Anthephorineae.

**Figure 1.**
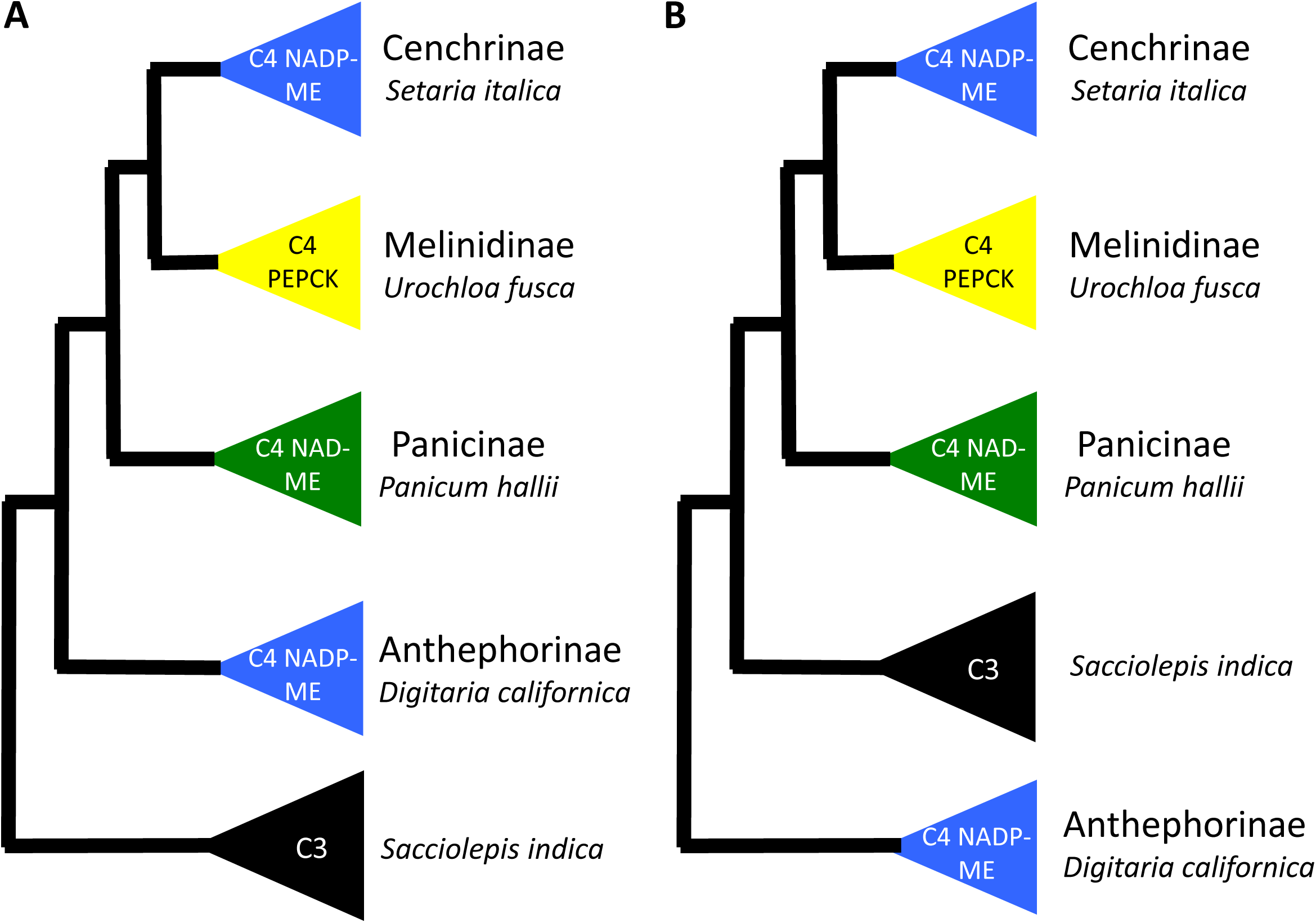
Phylogenetic relationships between a subset of species in the grass tribe Paniceae (Poaceae). The photosynthetic type (C_3_ or C_4_) and C_4_ sub-type of each species is labeled in the colored triangle next to it. *NADP-ME* = *NADP-DEPENDENT MALIC ENZYME*, *PCK* = *PHOSPHONENOLPYRUVATE CARBOXYKINASE*, *NAD-ME* = *NAD-DEPENDENT MALIC ENZYME*. A) Phylogeny based on nuclear genes (Vicentini et al. 2008 and Washburn et al. 2017). B) Phylogeny based on chloroplast genes (Washburn et al. 2015).

How and why C_4_ photosynthesis and its sub-types evolved has been investigated for many years (Raghavendra, 1980; Rawsthorne, 1992; Sage, 2001; Sage, 2004; Langdale, 2011; Sage et al., 2012; Schluter and Weber, 2020). Current hypotheses suggest an intermediate C_3_-C_4_ stage in which a photorespiratory pump operated (Sage, 2004; Sage et al., 2012; Heckmann et al., 2013; Mallmann et al., 2014; Bräutigam and Gowik, 2016; Blätke and Bräutigam, 2019). Each C_4_ sub-type would require at least some distinct evolutionary innovations, and the question of how or why multiple sub-types would evolve from the same C_3_ ancestor remains unanswered, although some evidence suggests it could be related to light quality and/or nutrient availability (Pinto et al., 2016; Sonawane et al., 2018; Blätke and Bräutigam, 2019; Arp et al., 2021).

To better determine whether the PEPCK pathway represents a true biochemical sub-type and investigate the extent to which C_4_ species make use of mixtures of C_4_ acid decarboxylases, global patterns of mRNA abundance were assessed from BS and M enriched samples across phylogenetically spaced C_4_ plants that were traditionally defined as exclusively using one of each of the C_4_ sub-types. These species belong to each subtribe of the CMP(A) described above. The C_3_ species *Sacciolepis indica*, another member of the Paniceae and sister to the CMP(A) clade (sister to CMP in chloroplast phylogeny and sister to CMPA in nuclear phylogeny), was included in the analysis to provide insight into the ancestral state and evolutionary transition from C_3_ to different C_4_ sub-types (Washburn et al., 2015; Washburn et al., 2017). A simple method was developed for isolating bundle sheath cells from the C_3_ species *Sacciolepis indica*.

We find that at least one species in the tribe appears to use PEPCK decarboxylation exclusively or nearly so, while the other species examined appear to be of mix subtype. Analysis of the C_3_ species *S. indica* shows low levels of C_4_ transcripts and an amenability to mechanical bundle sheath separation procedures not previously seen in C_3_ species. These observations lead us to hypothesize that S. indica may lie somewhere on the spectrum of C_3_-C_4_ intermediates or represent a reversion from an ancestral C_3_-C_4_ intermediate.

## Results

### M And BS Extraction And Distribution Of Transcripts Encoding The Core C_4_ Cycle

Four C_4_ species from the Paniceae tribe were chosen to represent the CMP(A) subtribes in the Paniceae (Figure 1). *Setaria italica* for Cenchrinae (NADP-ME), *Urochola fusca* for Melinidinae (PEPCK), *Panicum hallii for* Panicinae (NAD-ME), and *Digitaria californica* for Anthephorineae (NADP-ME). *Sacciolepis indica* was chosen to represent the closest C_3_ relative to the group.

Microscopic examination of leaves of *S. italica, U. fusca, P. hallii* and *D. californica*, from which M cell contents had been extracted, showed bands of cells containing low chlorophyll content (Figure 2A-D) a phenotype consistent with efficient removal of M content (Covshoff et al., 2013; John et al., 2014). In addition, after mechanical isolation of leaves, BS preparations of high purity for all C_4_ species were generated (Figure 2A-D). Separation of BS strands was also successful for the C_3_ species *Sacciolepis indica,* something that to our knowledge has not been successful in any other C_3_ species (Figure 2E). Analysis of transcripts derived from core C_4_ genes showed clear differences in abundance between M and BS samples from the C_4_ species. For example, transcripts derived from *CARBONIC ANHYDRASE* (*CA*), *PHOSPHOENOLPYRUVATE CARBOXYLASE* (*PEPC*) and *PYRUVATE, ORTHOPHOSPHATE DIKINASE* (*PPDK*) genes preferentially accumulated in M cells (Figure 3A). In contrast, transcripts derived from the *RUBISCO SMALL SUBUNIT* (*RBCS*) and *RUBISCO ACTIVASE* (*RCA*) as well as either *NADP-ME*, *NAD-ME* or *PEPCK* were more abundant in BS strands (Figure 3B). The abundance of transcripts relating to C_4_ photosynthesis in the C_3_ species *S. indica* were also consistent with current knowledge of metabolism in the BS of C_3_ species. For example, RBCS and RCA were more abundant in whole leaf samples than in the BS.

**Figure 2.**
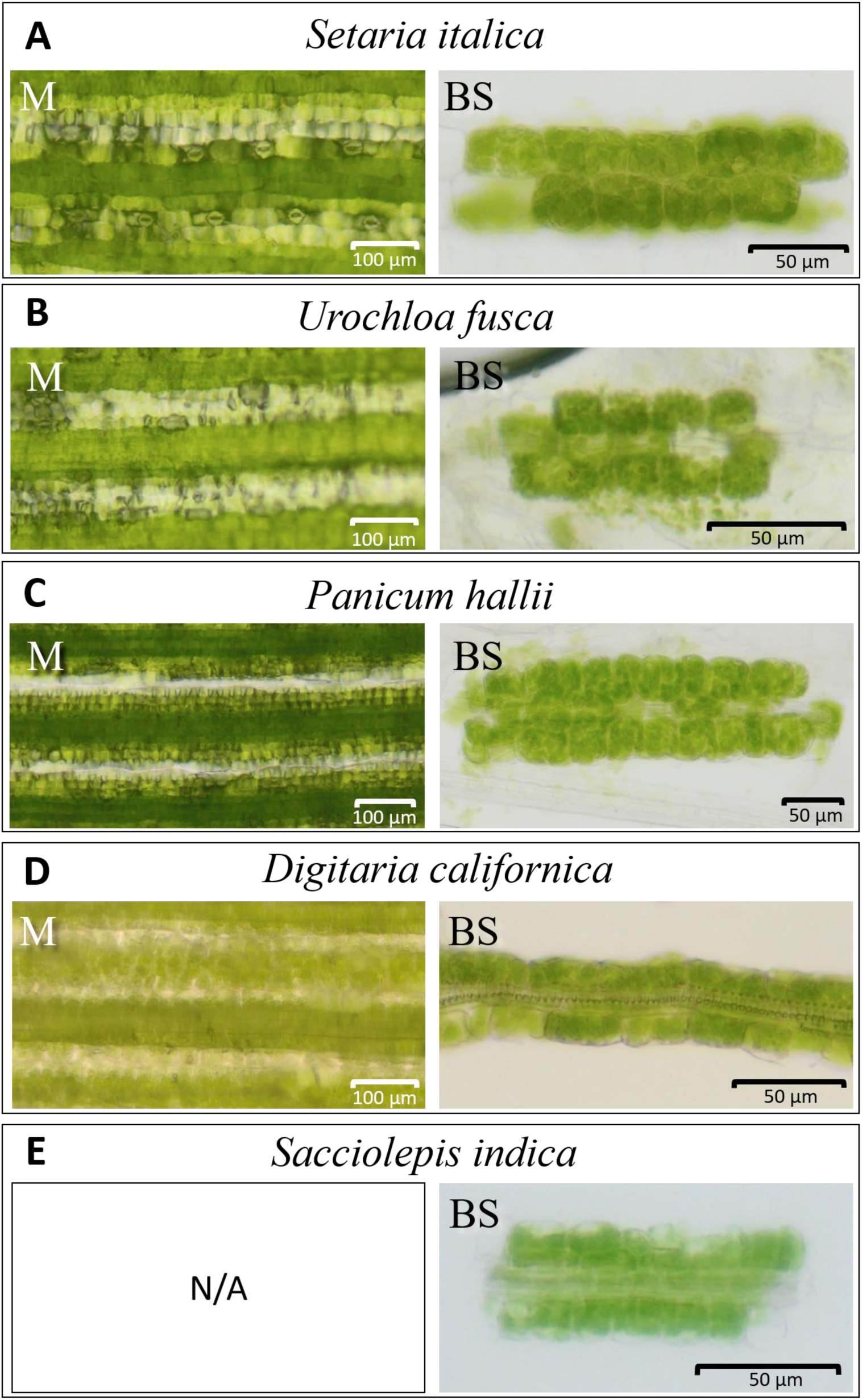
Representative whole leaf and bundle strands. Images from leaves that have been rolled to remove mesophyll (M) contents or bundle sheath (BS) strands after isolation. A) *Setaria italic*a, B) *Urochloa fusca*, C) *Panicum hallii*, D) *Digitaria californica*, E) *Sacciolepis indica*. All species use the C_4_ pathway except E, *Sacciolepis indica* which is a C_3_ plant. The bands of cells with low chlorophyll content in M images represent the position of mesophyll cells that have collapsed and had their contents expelled during the rolling procedure. Scale bars are depicted.

**Figure 3.**
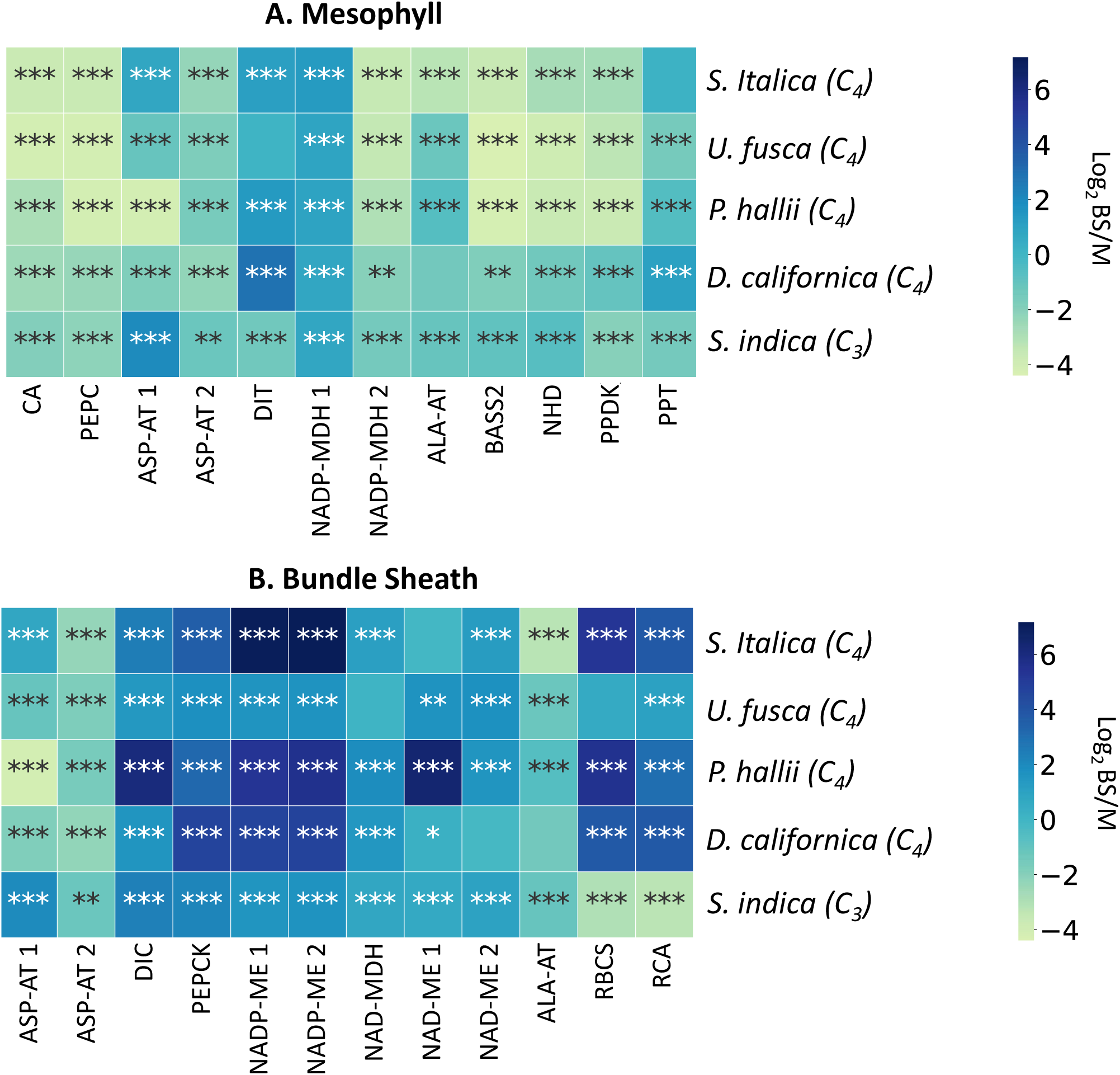
Log_2_ fold change between mesophyll (M) and bundle sheath (BS) enriched mRNA transcripts. Species used are *Setaria italica, Urochloa fusca, Panicum hallii* and *Digitaria californica*, and *Sacciolepis indica.* Note that for *S. indica*, a C_3_ species, whole leaf data is used in place of M. Genes depicted encode proteins of the core C_4_ cycle that are known to be preferentially expressed in either: A) M, or B) BS cells. The number of asterisks in each box represents the p-value. *** p < 0.001, ** p <0.01, * p < 0.05. *CA* = *CARBONIC ANHYDRASE*, *PEPC* = *PHOSPHOENOLPYRUVATE CARBOXYLASE*, *NADP-MDH* = *NADP-DEPENDENT MALATE DEHYDROGENASE, PPDK* = *PYRUVATE,ORTHOPHOSPHATE DIKINASE*, *NADP-ME* = *NADP-DEPENDENT MALIC ENZYME*, *PEPCK* = *PHOSPHONENOLPYRUVATE CARBOXYKINASE*, *NAD-ME* = *NAD-DEPENDENT MALIC ENZYME*, *NAD-MDH* = *NADP-DEPENDENT MALATE DEHYDROGENASE*, *RCA* = *RUBISCO ACTIVASE*, *RBCS* = *RUBISCO SMALL SUBUNIT*, *ASP-AT* = *ASPARAGINE-AMINOTRANSFERASE,* and *ALA-AT* = *ALANINE-AMINO TRANSFERASE*. DIT = *DICARBOXYLATE TRANSPORTER 1*, BASS2 = *SODIUM BILE ACID SYMPORTER 2*, NHD = *SODIUM:HYDROGEN ANTIPORTER*, DIC = *MITOCHONDRIAL DICARBOXYLATE CARRIER, PPT = PHOSPHATE/PHOSPHOENOLPYRUVATE TRANSLOCATOR.* The addition of a space and a number after the enzyme name indicates that multiple genes where mapped that may perform this function.

### Some Paniceae Lineages Use Classical Sub-Types and Others Mix C_4_ Acid Decarboxylases

*Setaria italica,* classically considered an NADP-ME sub-type species, showed high transcript levels for *NADP-ME* and *NADP-MDH* in BS and M cells respectively (Figure 4A). In addition, consistent with the *NADP-ME* sub-type, in BS strands of *S. italica* transcripts encoding *PEPCK*, *NAD-ME*, *NAD-MDH*, *ASP-AT*, and *ALA-AT* were detected at low levels. Enzyme activity assays also indicated high levels of NADP-ME in *S. italica* (Figures 5 and 6). Surprisingly high levels of PEPCK enzyme activity were also found in *S. italica,* but this was not the case for *PEPCK* transcripts. This may be explainable by the differences in growth chamber conditions, though slight, between RNA samples and enzyme activity samples or an alternate protein may have generated this activity.

**Figure 4.**
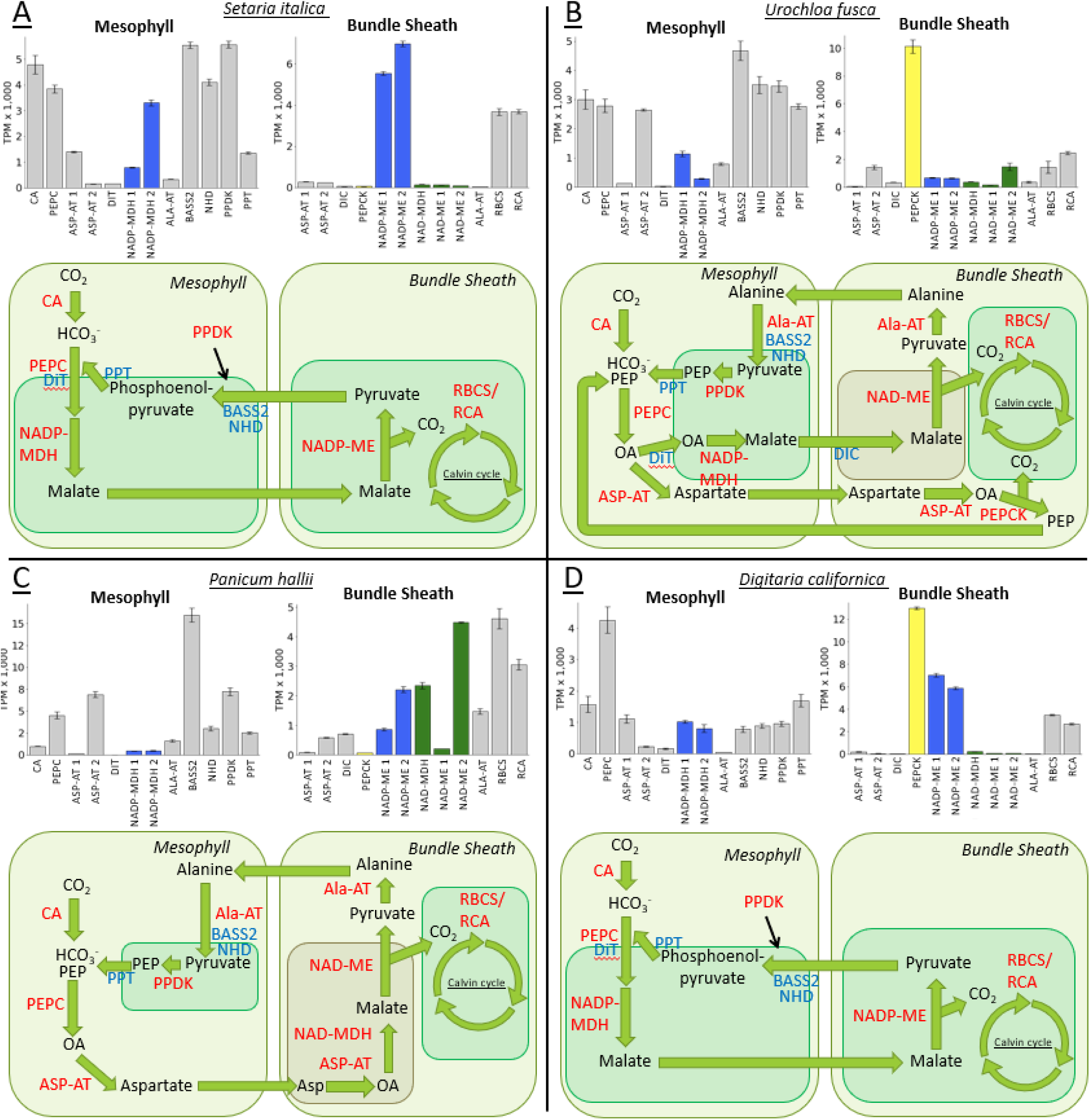
Relative transcript abundance between core C_4_ enzymes within mesophyll (M) and bundle sheath (BS) extracts. Data is displayed for: A) *Setaria italica,* B) *Urochloa fusca,* C) *Panicum hallii,* and D) *Digitaria californica*. The schematics below each histogram indicate the enzyme complement associated with each of the three biochemical sub-types. *CA* = *CARBONIC ANHYDRASE*, *PEPC* = *PHOSPHOENOLPYRUVATE CARBOXYLASE*, *NADP-MDH* = *NADP-DEPENDENT MALATE DEHYDROGENASE, PPDK* = *PYRUVATE,ORTHOPHOSPHATE DIKINASE*, *NADP-ME* = *NADP-DEPENDENT MALIC ENZYME*, *PEPCK* = *PHOSPHONENOLPYRUVATE CARBOXYKINASE*, *NAD-ME* = *NAD-DEPENDENT MALIC ENZYME*, *NAD-MDH* = *NADP-DEPENDENT MALATE DEHYDROGENASE*, *RCA* = *RUBISCO ACTIVASE*, *RBCS* = *RUBISCO SMALL SUBUNIT*, *ASP-AT* = *ASPARAGINE-AMINOTRANSFERASE,* and *ALA-AT* = *ALANINE-AMINO TRANSFERASE*. DIT = *DICARBOXYLATE TRANSPORTER 1*, BASS2 = *SODIUM BILE ACID SYMPORTER 2*, NHD = *SODIUM:HYDROGEN ANTIPORTER*, DIC = *MITOCHONDRIAL DICARBOXYLATE CARRIER, PPT = PHOSPHATE/PHOSPHOENOLPYRUVATE TRANSLOCATOR.* The addition of a space and a number after the enzyme name indicates that multiple genes where mapped that may perform this function. Error bars are plus or minus the standard error across replicates.

**Figure 5.**
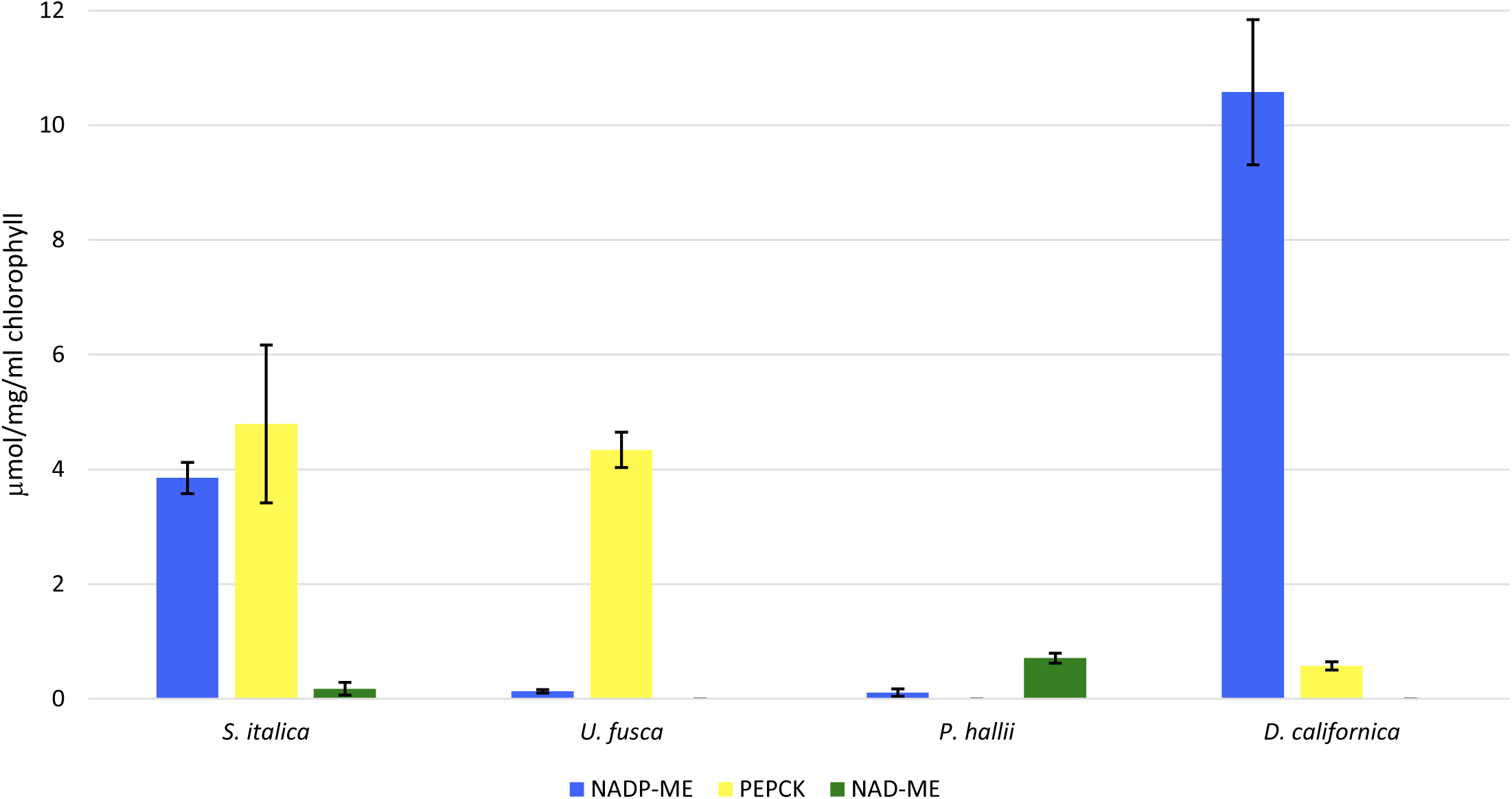
Enzyme activities of C_4_ decarboxylases. NADP-DEPENDENT MALIC ENZYME (NADP-ME), PHOSPHOENOLPYRUVATE CARBOXYKINASE (PEPCK), and NAD-DEPENDENT MALIC ENZYME (NAD-ME) for *Setaria italica, Urochloa fusca, Panicum hallii,* and *Digitaria californica*. Error bars represent plus or minus the standard error across replicates.

**Figure 6.**
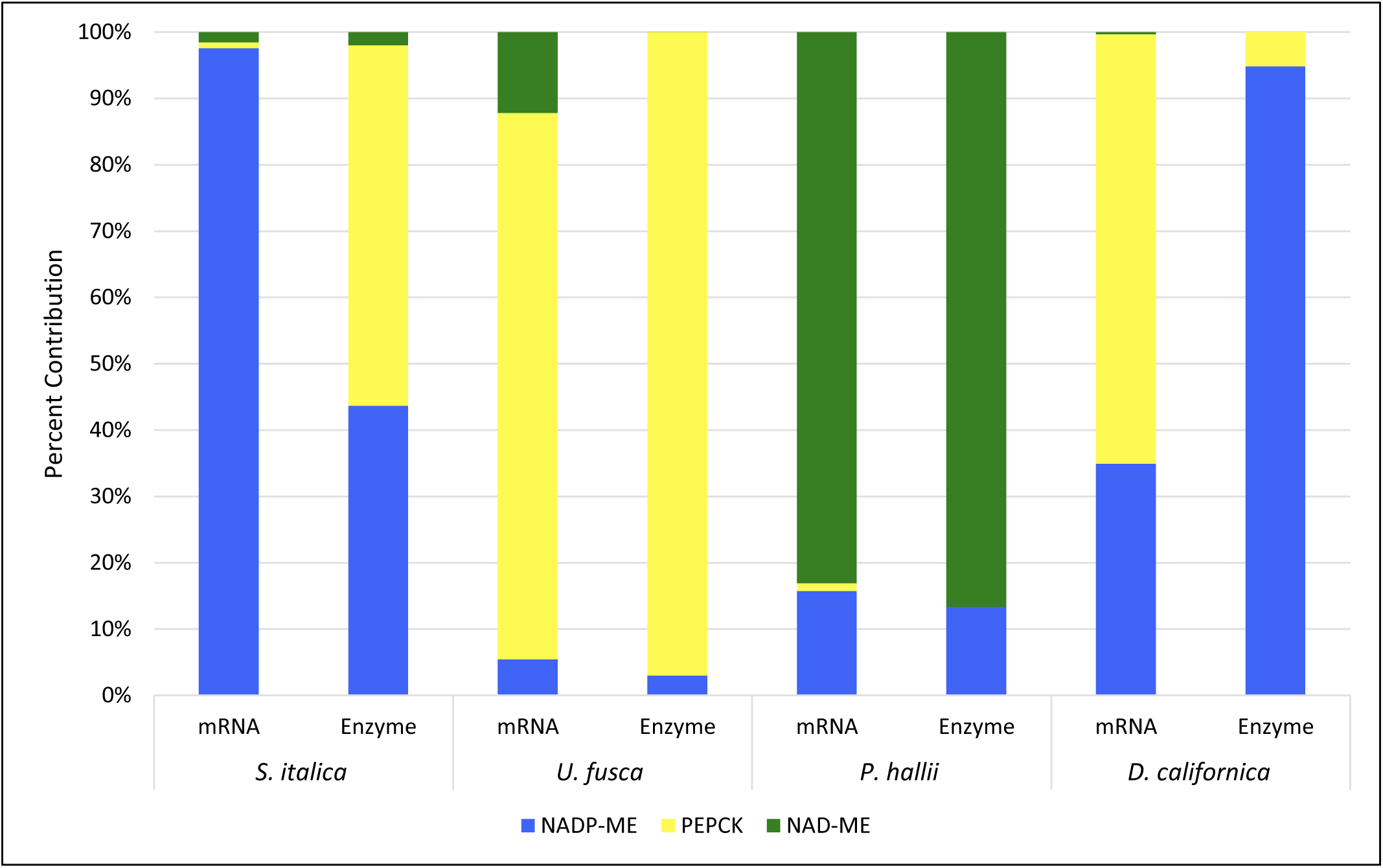
Relative transcript accumulation and enzyme activities of C_4_ decarboxylases. *NADP-DEPENDENT MALIC ENZYME* (*NADP-ME*), *PHOSPHOENOLPYRUVATE CARBOXYKINASE* (*PEPCK*), and *NAD-DEPENDENT MALIC ENZYME* (*NAD-ME*) transcript accumulation and enzyme activities for *Setaria italica, Urochloa fusca, Panicum hallii,* and *Digitaria californica*. Values are represented as a percentage of the total of all three decarboxylase values. Enzyme and mRNA data were not collected at the same time, but under closely matching environmental conditions.

*Urochola fusca* is classically thought to exclusively use *PEPCK* to release CO_2_ in the BS. The patterns of transcript accumulation in M and BS strands of *U. fusca* are consistent with *PEPCK* functioning in this species with very little to no supplemental decarboxylation from either *NADP-ME* or *NAD-ME* (Figure 4B). BS stands contained barely detectable levels of transcripts encoding *NADP-ME* and *NAD-ME*, but very high levels of those encoding *PEPCK*. Enzyme activity in *U. fusca* was consistent with transcript abundances, PEPCK having high levels and the other two decarboxylases very low levels (Figures 5 and 6). In addition, consistent with the cycling of aspartate and alanine between the two cell-types, transcripts derived from genes encoding both *ASP-AT* and *ALA-AT* were detectable in the two cell-types (Figure 4B).

In contrast to the above analysis of *S. italica* and *U. fusca* which seem to fit with an exclusive use of one decarboxylation enzyme, analysis of *P. hallii* and *D. californica* indicated they potentially use multiple C_4_ acid decarboxylases during photosynthesis (Figure 4C-D). Although *P. hallii* is classically considered to use *NAD-ME* in addition to high levels of transcripts encoding *NAD-ME*, *NAD-MDH*, *ASP-AT* and *ALA-AT*, unexpectedly high levels of transcripts encoding *NADP-ME* were detected in the BS (Figure 4C). *NAD-ME* transcript levels where still more than twice those of *NADP-ME*, and the enzyme assays found much higher relative levels of NAD-ME than NADP-ME (Figures 5 and 6). In the case of *Digitaria californica* which is thought to belong to the *NADP-ME* sub-type, although transcripts encoding *NADP-ME* and *NADP-MDH* were abundant in BS and M samples respectively, *PEPCK* levels were more than double those of *NADP-ME* in the BS (or perhaps the levels are similar if one considers the possibility that both *NADP-ME* genes here mapped are resulting in similar functional products). Enzyme assay results showed high levels of NADP-ME and much lower levels of PEPCK consistent with the traditional subtype classification of this species (Figures 5 and 6).

To further confirm or refute these findings RNA *in situ* hybridizations were undertaken. Transcript accumulation by *in situ* hybridization experiments were consistent in signal strength with the RNA-seq results from above and indicated that the signals clearly localized to the expected anatomical locations (Supplemental Figure 1). Previous enzyme assay results for species closely related to the ones sampled here are also extremely consistent with the RNA-Seq results (Supplemental Figure 2). It should be noted that the conditions for mRNA, *in situ*, and enzyme assay sample collection were not identical (see Materials and Methods and Discussion sections).

### C_4_ Pathway Transporters

The transcript abundance of various transporters related to C_4_ photosynthesis were examined. Some transporters had low or undetectable levels such as *DICARBOXYLATE TRANSPORTER 1 (DIT1).* Others, such *SODIUM BILE ACID SYMPORTER 2 (BASS2)*, *SODIUM:HYDROGEN ANTIPORTER (NHD)*, *MITOCHONDRIAL DICARBOXYLATE CARRIER (DIC),* and *PHOSPHATE/PHOSPHOENOLPYRUVATE TRANSLOCATOR (PPT)*, had variable levels of transcript abundance across the species.

### C_4_ Transcript Abundance Levels In The Closest C_3_ Relative To The MPC(A) Clade

Transcript abundance levels from *S. indica*, a C_3_ species that is part of the most closely related out group to the CMP(A) clade, were generally consistent with expectations for a C_3_ species (Figures 3 and 7). RBCS and RCA are more highly expressed in whole leaf tissue than in BS extracts. Transcripts related to C_4_ photosynthesis are also expressed at a low level in both whole leaf and BS.

**Figure 7.**
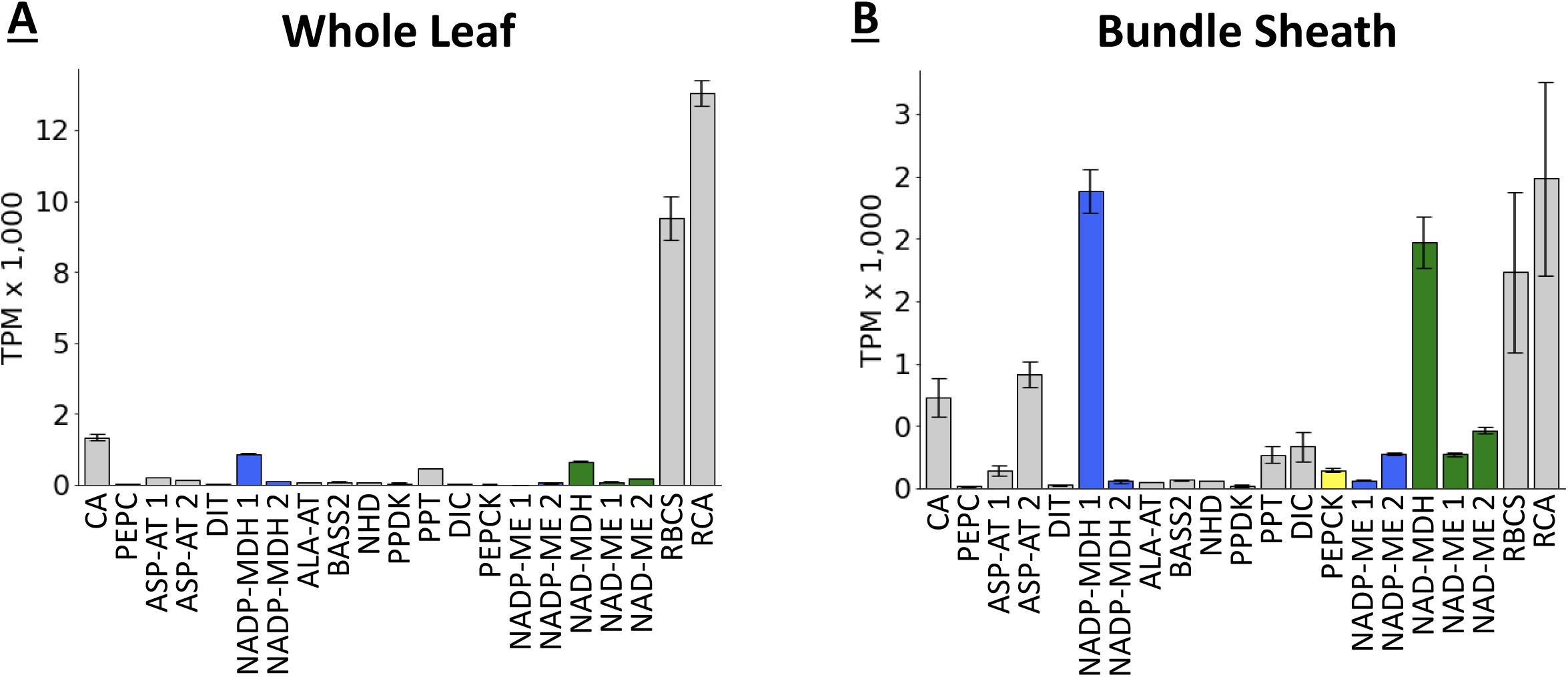
Relative transcript abundance between core C_4_ enzymes within whole leaf and bundle sheath (BS) extracts from *Sacciolipis indica*. A) Whole leaf, B) Bundle sheath*. CA* = *CARBONIC ANHYDRASE*, *PEPC* = *PHOSPHOENOLPYRUVATE CARBOXYLASE*, *NADP-MDH* = *NADP-DEPENDENT MALATE DEHYDROGENASE, PPDK* = *PYRUVATE,ORTHOPHOSPHATE DIKINASE*, *NADP-ME* = *NADP-DEPENDENT MALIC ENZYME*, *PEPCK* = *PHOSPHONENOLPYRUVATE CARBOXYKINASE*, *NAD-ME* = *NAD-DEPENDENT MALIC ENZYME*, *NAD-MDH* = *NADP-DEPENDENT MALATE DEHYDROGENASE*, *RCA* = *RUBISCO ACTIVASE*, *RBCS* = *RUBISCO SMALL SUBUNIT*, *ASP-AT* = *ASPARAGINE-AMINOTRANSFERASE,* and *ALA-AT* = *ALANINE-AMINO TRANSFERASE*. DIT = *DICARBOXYLATE TRANSPORTER 1*, BASS2 = *SODIUM BILE ACID SYMPORTER 2*, NHD = *SODIUM:HYDROGEN ANTIPORTER*, DIC = *MITOCHONDRIAL DICARBOXYLATE CARRIER, PPT = PHOSPHATE/PHOSPHOENOLPYRUVATE TRANSLOCATOR.* The addition of a space and a number after the enzyme name indicates that multiple genes where mapped that may perform this function. Error bars are plus or minus the standard error across replicates.

Comparisons between the *S. indica* (C_3_) BS enriched and whole leaf samples showed significant (adjusted p-value < 0.001) BS over-abundance for 912 gene models and whole leaf (WL) over-abundance for 746 gene models (Supplemental Table 1). Of these over-abundant genes, significant Gene Ontology (GO) enrichment (adjusted p-value < 0.05) was found for 16 different GO terms for the BS, and 14 GO terms for the M (Supplemental Table 1). The different GO terms enriched in BS cells related to diverse processes including cellular transport, molecular binding, and cell wall and membrane components. The WL GO terms related to photosystems I & II, photosynthesis, and other processes (Supplemental Table 1).

### Gene Expression in C_3_ versus C_4_ Bundle Sheath Cells

The experimental design provides the opportunity to compare the BS expression from a C_3_ species against BS expression in four closely related C_4_ species. A total of 357 gene models displayed significantly (padj < 0.001, log2 fold change > 2) higher transcript abundance levels over the C_3_ BS in all four C_4_ species in the analysis (Supplemental Table 2). Many of these genes are expected, such as photosystem I and II subunits, cytochrome b_6_f, cyclic electron chain proteins, Calvin cycle proteins, cellulose synthase, *Pectinacetylesterase*, starch synthase, and others. The remaining genes are potential candidates involved in C_4_ photosynthesis that deserve further molecular and biochemical investigation (Supplemental Table 2).

## Discussion

### The PEPCK Sub-Type

The dominance of *PEPCK* transcript and enzyme activity over *NADP-ME* and *NAD-ME* in *U. fusca* provides evidence for the biological relevance of the classical PEPCK sub-type (Figures 4-6, Supplemental Figure 2). These data contrast with proposals that PEPCK cannot function on its own but rather is always ancillary to one of the other two C_4_ acid decarboxylases (Furbank, 2011; Bräutigam et al., 2014; Wang et al., 2014). While this notion may be true in some cases, the current results suggest that it is not the case for *U. fusca*. Furthermore, our findings are supported by earlier measurements of enzyme activity made within the sub-tribe Melinidinae, where PEPCK was also shown to be highly dominant over the other sub-types (Gutierrez et al., 1974; Prendergast et al., 1987; Lin et al., 1993), and also indicate that these differences in activity are due to differences in steady-state transcript abundance rather than post-transcriptional modifications that act to suppress accumulation of NADP-ME and NAD-ME.

### C_4_ Sub-Type Mixing

Of the four C_4_ species examined, *P. hallii* and *D. californica* show the most evidence of sub-type mixing. The potential for mixing has previously been considered in *Panicum virgatum*, a close relative of *P. hallii* (Zhang et al., 2013; Meyer et al., 2014; Rao and Dixon, 2016; Rao et al., 2016). Rao et al. (2016) suggested that the high abundance of *NADP-ME* transcripts may be accounted for by post-transcriptional or translational modifications but experimental evidence for testing that hypothesis is lacking.

*Digitaria californica* also showed some evidence of sub-type mixing. In this case, *NADP-ME* and *PEPCK* transcripts were both reasonably abundant. Although this species is classically considered to belong to the NADP-ME sub-type, transcripts encoding *PEPCK* were more than double the abundance of those of *NADP-ME*. *ASP-AT* transcript levels, which are also associated with the *PEPCK* sub-type, were high as well. However, enzyme activity data does not support this idea with NADP-ME having much higher levels than PEPCK. Some of these differences between transcript and enzyme levels could be due the enzyme activity assays being carried out on samples from a different growth chamber with somewhat different conditions than the RNA sequencing samples (See Materials and Methods).

### *Sacciolepis indica* and the C_3_ Bundle Sheath

Analysis of transcript abundance in M and BS cells from C_3_ species that are closely related to C_4_ species is critical to understanding how C_4_ photosynthesis evolved and how it can be engineered for enhanced crop production. Although transcripts loaded onto ribosomes in the BS of C_3_ *Arabidopsis thaliana* have been assessed, and this analysis provided insight into the role of the BS in eudicot plants (Aubry et al., 2014), to our knowledge, there are no equivalent data from a monocotyledonous lineage in which both C_3_ and C_4_ species are found. The ability to mechanically isolate intact BS cells indicates that *S. indica* has very strong BS cell walls, similar to those found in C_4_ species. However, all other currently available data including phylogenetic placement and RNA-seq from this study are consistent with *S. indica* being a C_3_ species. The relatively high levels of C_4_ related transcripts in the BS of *S. indica* are consistent with previous work on cells around the veins of C_3_ plants (Hibberd and Quick, 2002; Brown et al., 2010; Shen et al., 2016). Together, these data support the concept that some C_3_ species are pre-adapted to adopt the C_4_ mechanism (Gould, 1989; Christin et al., 2009; Brown et al., 2011; Christin et al., 2015; Washburn et al., 2016; Williams et al., 2016; Reyna-Llorens et al., 2018; Burgess et al., 2019). Another interesting hypothesis from this study is that perhaps *S. indica* represents a step on the pathway to becoming a C_3_-C_4_ intermediate, or maybe it represents a reversion to C_3_ photosynthesis from an ancestral C_3_-C_4_ intermediate (Sage, 2004; Bräutigam and Gowik, 2016).

### The *S. indica* C_3_ BS shows marked similarities and differences to the BS in other species

Aubry and colleagues (2014) investigated the functions of *Arabidopsis thaliana* BS cells by labeling ribosomes within the cell type and sequencing the mRNA resident in the ribosomes. In general, our *S. indica* BS cells displayed similar patterns to those seen in Arabidopsis. Of the 912 significantly over-abundant gene models in the *S. indica* BS, 50 of them have Arabidopsis homologs that were significantly over-abundant in BS cells within the Aubry study (Aubry et al., 2014). These genes have annotated functions relating to transport (nucleotide, peptide, amino acid, sulphate, metals, ABS transporters), metal handling, transcription regulation, protein degradation, cell wall modification, amino acid metabolism, hormone metabolism, and ATP synthesis. Other functional annotations present both in the Arabidopsis and *S. indica* upregulated BS gene sets (but not necessarily from homologous genes) included: nitrogen metabolism, glutamine synthetase, tryptophan, ethylene induced signaling and regulation, lipid metabolism, trehalose metabolism, phenylpropanoid metabolism, and sulfur regulation (Supplemental Table 1).

Similarly to Arabidopsis and maize, several trehalose metabolism genes were found to be overexpressed within the *S. indica* BS, supporting the hypothesis that metabolism of trehalose is an ancestral BS function (Chang et al., 2012; Aubry et al., 2014). The data are also consistent with the hypothesis that BS cells play an important role in sulfur transport and metabolism and in nitrogen metabolism (Aubry et al., 2014). We do note, however, that some sulfur metabolism related genes shown to be enriched in Arabidopsis BS were actually found to be depleted in the *S. indica* BS. Overall, the *S. indica* BS samples are highly consistent with previous studies on Arabidopsis and rice BS, indicating both the validity of the mechanical C_3_ BS isolation done here, and the conservation of C_3_ functions across these divergent species (Aubry et al., 2014; Hua et al., 2021).

### The Evolution of Three C4 Sub-types in the MPC(A)

For the majority of C_4_ genes examined, all five species appear to use orthologous genes, or at least their transcripts mapped to the same *S. bicolor* gene (Figures 4 and 7). This is based on the common assumption that the highest expressed gene in each species/tissue is the one being used. For CA, PEPC, PPDK, PPT, NHD, ALA-AT, NAD-MDH, PEPCK, and NAD-ME the highest expressed gene in all species was clearly the same, although in some species the gene expression was so low it is likely non-functional. NADP-MDH, NADP-ME, and ASP-AT are less clear with the highest gene being different between some species but also often having high abundance levels for both genes making it hard to conclude that the lower gene is not relevant. The lack of good genomic resources for all species involved makes it difficult to conclude if the same genes are in fact being used by all species, and therefore potentially the result of a single recruitment, or if the genes are simply close homologs and recruited to C_4_ separately in different lineages.

Ancestral state reconstructions were also performed based on the transcript abundance and enzyme activity data, however, these analyses were inconclusive and have been excluded due to the low phylogenetic sampling of the clade within this study.

## Materials and Methods

### Plant Materials

Accessions from five plant species were used in this study: *Setaria italica* yugu1, *Urochloa fusca* LBJWC-52, *Panicum hallii* FIL2, *Digitaria californica* PI 364670, and *Sacciolepis indica* PI 338609. More details on the accessions can be found in Washburn et al. (2015) with exception of *P. hallii* FIL2, obtained from Thomas Juenger of the University of Texas at Austin with further details at *Panicum hallii* v2.0, DOE-JGI, http://phytozome.jgi.doe.gov/.

Plant materials for RNA sequencing were grown in controlled growth chambers at the University of Missouri in Columbia. Plants were grown under 16 hours of light (from 6:00-20:00) and 8 hours of darkness with temperatures of 23C during the day and 20C at night. Lights were placed between 86-88 cm above the plants. Plantings were grown in 4 replicates in a completely randomized design with 32 plants per replicate (except for *Sacciolepis indica* where plants were smaller and grown with 64 plants per replicate). The third leaf was sampled between 11:00 and 15:00 using established leaf rolling and mechanical BS isolation methods with some modifications (See Supplemental Protocol 1) (Sheen and Bogorad, 1985; Chang et al., 2012; Covshoff et al., 2013; John et al., 2014). Due two time and cost constraints only 3 of the 4 replicates (each based on a pool of plants) were processed for sequencing.

The protocol used for obtaining BS strands in *S. indica* was the same as that used for the C_4_ species. Variations on the amount of time for each blending step were investigated, but only resulted in higher levels of contamination as viewed under a microscope. That said, even when microscopic examination indicated higher contamination levels in some *S. indica* BS samples, transcript abundance levels were qualitatively similar to samples with less apparent contamination. One step that may have been key to the isolation of C_3_ BS strands, was the use of leaf rolling on the sampled leaves just prior to the BS strand isolation procedure (Furbank et al., 1985; John et al., 2014). This enables the removal of at least some M sap prior to BS isolation.

M enriched samples where not successfully obtained for *S. indica* in this study because of the sensitivity of the C_3_ leaves to rolling. Very small amounts of leaf rolling pressure resulted in the leaves becoming damaged to the point that the purity of M sap obtained from them was called into question. Leaves that were rolled gently enough not to damage the BS strands and contaminate the M sap resulted in sap with insufficient quantities of RNA for sequencing. It is our opinion that M sap could be sampled using: 1) low input mRNA extraction and sequencing procedures, 2) a more precise instrument for leaf rolling such as that described by Leegood (1985), and/or 3) further experimentation with the developmental stage at which M sap is extracted.

This resulted in 3 replicates of BS and M (or whole leaf tissue for the C_3_) for each of the 5 species used for a total of 30 samples used in RNA extraction, sequencing, and analysis.

### Sequencing

RNA was extracted using the PureLink® RNA Mini Kit (Invitrogen, Carlsbad, CA, USA) and mRNA-seq libraries were constructed and sequenced by the University of Missouri DNA Core Facility using the TruSeq Stranded mRNA Sample Prep Kit (Illumina, Inc., San Diego, CA, USA) and the Illumina HiSeq and NextSeq platforms.

### Analysis

Each mRNA sample was quality trimmed and mapped to the *Sorghum bicolor* genome version 3.1.1 (Paterson et al., 2009; DOE-JGI, 2017). All species were mapped to *S. bicolor* because reference genomes do not exist for some of the species in this study and the reference genomes that do exist are of variable quality leading to bias. Mapping all species to S. bicolor, which is equally related to all, also allows for orthology assignment during the mapping step as opposed to later in the process. Raw sequence was processed using Trimmomatic and Trinity v2.8.4 following the workflows outlined on the Trinity website (Grabherr et al., 2011; Haas et al., 2013; Bolger et al., 2014). This processing included the use of eXpress and Bowtie2 for read mapping and counting as well as edgeR and DESeq2 for differential expression analysis (Robinson et al., 2010; Li and Dewey, 2011; Langmead and Salzberg, 2012; McCarthy et al., 2012; Love et al., 2014). A list of known C_4_ photosynthesis genes was compiled based on the literature; a custom script and BLAST were then used to find the appropriate homologous genes and to compare their relative abundance levels (Camacho et al., 2009; Chang et al., 2012; Covshoff et al., 2013; Bräutigam et al., 2014; John et al., 2014; Tausta et al., 2014; Rao et al., 2016). For comparisons across all cell types and species within this study, the Trimmed Mean of M-values (TMM) method described by Robinson and Oshlack (2010) as implemented in DESeq2 was used. All scripts and workflows used in the analysis can be found in a Bitbucket repository at https://bitbucket.org/washjake/paniceae_c4_m_bs_mrna/.

### Enzyme assays

Enzyme activity assays were performed based on methods described in (Ashton et al., 1990; Marshall et al., 2007). Samples were grown in growth chambers at the University of Cambridge, UK and growth conditions were matched as closely as possible to those above. The temperature was a constant 20C, 60% humidity, 300 µmol light, and 16 hours of light. Samples were prepared by grinding leaf tissue with a pestle and mortar in extraction buffer then centrifuged at 13000 x g and supernatant taken.

For PEPCK assays, we used the Walker and Leegood method that was developed to reduce proteolysis, and which has been used extensively (Walker et al., 1995; Häusler et al., 2001; Marshall et al., 2007; Sommer et al., 2012; Sharwood et al., 2016). As it has been previously reported that the forward (de-carboxylation) reaction is about 2.6 times faster than the reverse (carboxylation) reaction, the PEPCK activity *in vivo* may be higher than what we measured (Ashton et al., 1990). PEPCK extraction buffer consisted of 200 mM Bicine-KOH, pH 9.0, 20 mM MgCl2 and 5 mM DTT. NAD-ME extraction buffer consisted of 50 mM HEPES-NaOH pH 7.2, 50 mM Tricine, 2 mM MnCl2, 5 mM DTT, 0.25% w/v PVP- 40000 and 0.5% Triton X-100. NADP-ME extraction buffer consisted of 50 mM HEPES-KOH pH 8.3, 50 mM Tricine, 5 mM DTT and 0.1 mM EDTA.

For PEPCK activity, assay buffer contained of 80 mM MES-NaOH pH 6.7, 0.35 mM NADH, 5 mM DTT, 2 mM MnCl2, 2 mM ADP and 50 mM KHCO3. Background rates were measured for five minutes then 1.2 units of malate dehydrogenase was added and rates measured for a further five minutes. Assays were initiated with the addition of 2mM Phosphoenolpyruvate (PEP). For NAD-ME activity, assay buffer contained 25 mM HEPES-NaOH pH 7.2, 5 mM L-malic acid, 2 mM NAD, 5 mM DTT, 0.2 mM EDTA. Background rates were measured for five minutes then 24 mM MnCl2 and 0.1 mM coenzyme A were used to initiate the reaction. For NADP-ME activity, assay buffer contained 25 mM Tricine-KOH pH 8.3, 5 mM L-malic acid, 0.5 mM NADP, 0.1 mM EDTA. Background rates were measured for five minutes, and the assays were initiated with 2mM MgCl2.

All assays were performed in 96 well plates at 25°C in a CLARIOstar plus plate-reader (BMG labtech) in 200 μl reactions with absorbance at 340 nm measured every 60 s until steady states were reached. Rates were calculated as the rate of reaction from the initial slope of the reaction minus any observed background rate. Rates were normalized to both protein concentration, measured using the Qubit protein assay (Life Technologies), and chlorophyll concentration, extracted using 80% acetone and calculated as in Porra et al. (1989).

### RNA *in situ* hybridization

Each of the genotypes were grown in a climate-controlled growth chamber at 50% relative humidity in 16:8 light to dark cycles at 29.4° C and 23.9° C day and night temperature, respectively. These conditions were different from the original mRNA samples due to the logistics of growth chamber availability. However, since the results appear to support those from the RNAseq at a lower temperature it does not appear this temperature difference had a strong impact. Replicates of fully expanded leaf three were harvested from each genotype when plants were at vegetative stage 4 (V4) when the fourth leaf collar was visible. Along the longitudinal length of the leaf blade, mid-sections of blade tissue were dissected in 3.7% FAA at 4° C. Samples were vacuum infiltrated and fixed overnight at 4° C in 3.7% FAA. Leaf samples were dehydrated through a graded ethanol series (50%, 70%, 85%, 95%, 100%) with 3 changes in 100% ethanol; all changes were 1 hour each at 4° C except for the last 100% ethanol, which was overnight at 4° C. Samples were then passed through a graded HistoChoice® (Sigma-Aldrich) series (3:1, 1:1, 1:3 ethanol: HistoChoice®) with 3 changes in 100% HistoChoice®; all changes were 1 hour each at room temperature. Samples were then embedded in Paraplast®Plus (McCormick Scientific), sectioned to 10 mm, and hybridized as described previously (Strable and Satterlee, 2021). Two fragments for *PEPCK* consisted of 450 base pairs (bp) of the CDS (synthesized from JSC4-6 and JSC4-7 primers) and 456 bp of the 3’ end that included UTR (JSC4-4 and JSC4-5). Two fragments for *NADP-ME* consisted of 790 base pairs (bp) of the CDS (JSC4-8 and JSC4-9) and 286 bp of the 3’ end that included UTR (JSC4-10 and JSC4-11). Fragments were subcloned into pCR 4-TOPO (Invitrogen) and confirmed by Sanger sequencing. Antisense or sense strand digoxygenin-UTP labeled RNA was generated for *PEPCK* and *NADP-ME* using a DIG RNA labeling kit (Roche). For *PEPCK* hybridizations, equal amounts of the two probes for *PEPCK* were mixed prior to hybridization. Similarly, for *NADP-ME* hybridizations, equal amounts of the two probes for *NADP-ME* were mixed prior to hybridization. Primer sequences for RNA in situ probes are provided in Supplemental Table 3.

### Accession numbers

Sequence data are available on NCBI SRA (https://www.ncbi.nlm.nih.gov/sra) under accession number ….

## Supplemental data files

Supplemental Table 1. Differentially expressed genes between the whole leaf and bundle sheath of *Sacciolepis indica*.

Supplemental Table 2. Differentially expressed genes that are upregulated in all four C4 species bundle sheath cells and down regulated in *Sacciolepis indica* bundle sheath.

## Author Contributions

All Authors contributed to drafting and revising the manuscript. JDW, JCP, GCC, SC, and JMH conceived of the work and experimental design. JDW, SC, SSK, and JMB developed and performed the leaf rolling experiments. JDW performed the bioinformatic analysis. JS performed the RNA *in situ* hybridization experiments. PD and JMH designed and performed the enzyme assay experiments. JDW has agreed to serve as the author responsible for contact and communication.

## Acknowledgments

This work was supported by grants from the University of Missouri (Mizzou Advantage, MU Research Board, and Molecular Life Science Fellowships), and the U.S. National Science Foundation (Award no. 1501406, 1710618, 1710973). Additional support comes from the U.S. Department of Agriculture, Agricultural Research Service.

