## Supplemental Figures for "Distinct C_4_ Sub-Types and C_3_ Bundle Sheath Isolation In The Paniceae Grasses"

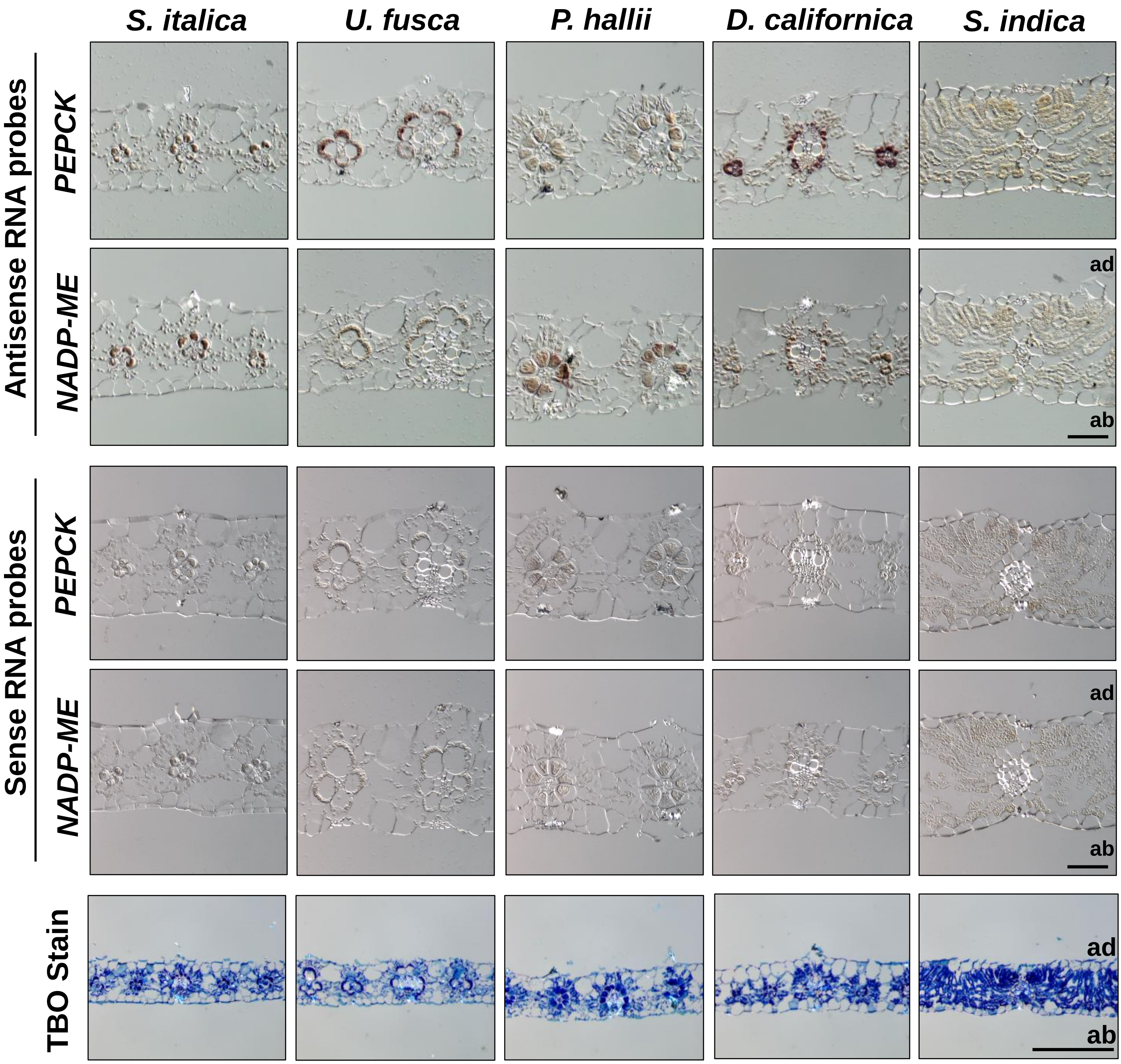


**Supplemental Figure 1. PHOSPHOENOLPYRUVATE *CARBOXYKINASE* (*PEPCK)* and *NADP-DEPENDENT MALIC ENZYME* (*NADP-ME*) transcript accumulation in fully expanded leaves.** Transverse sections through midpoints of fully expanded leaf 4 hybridized with antisense and sense RNA. Toluidine blue-O staining of fully expanded leaves. ad, adaxial; ab, abaxial. Scale bars, 200 µm.

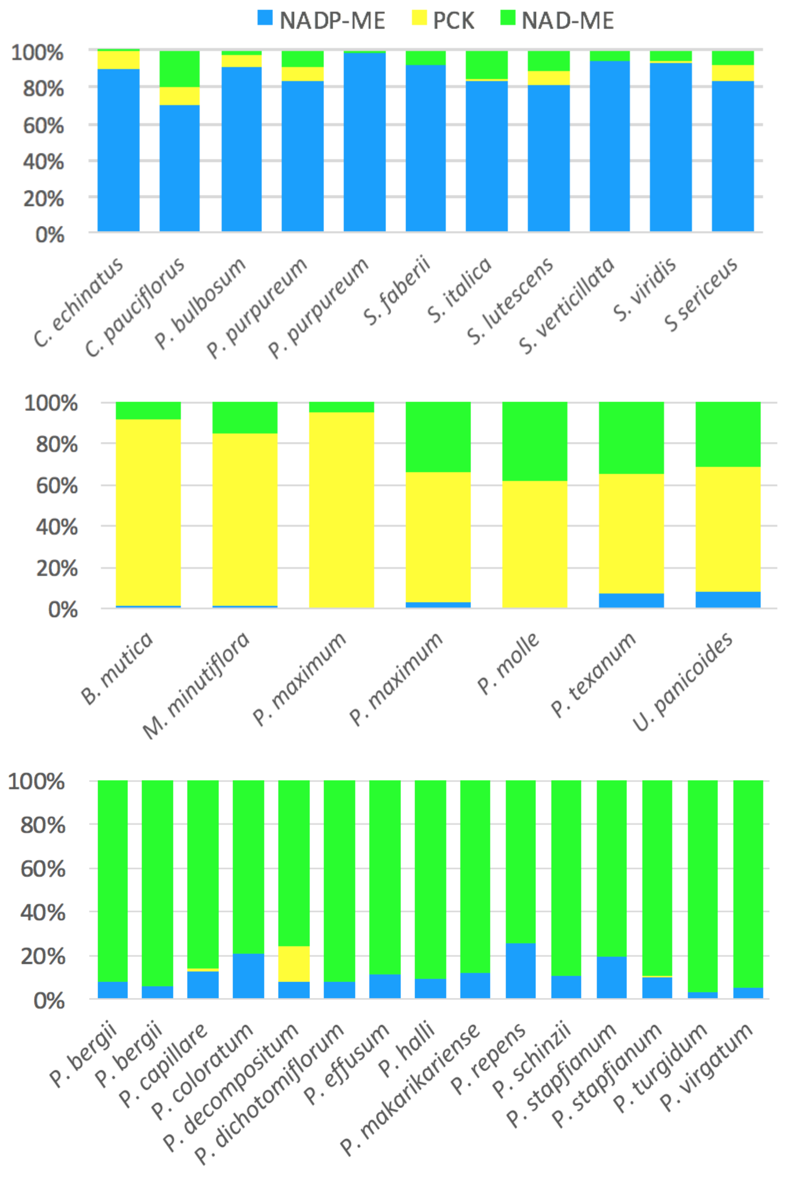


**Supplemental Figure 2. PCK, NADP-ME, and NAD-ME enzyme abundance levels in different Paniceae species.**
Data taken from Gutierrez et al., 1974. Planta 119:279-300, Prendergast et al., 1987. Funct. Plant Biol. 14:403-420, Lin et al., 1993. Funct. Plant Biol. 20:757-769.

**Supplemental Table 3. Primer sequences for *in situ* hybridization.**

| Primer ID | SEQUENCE | GENE | GENE ID | Probe size | Location |
|---|---|---|---|---|---|
| JSC4-4 | GAGAATCTCCTGAGGCTCGG | GRMZM2G001696 | PCK | 456 | 3' end |
| JSC4-5 | ACAGGGGGCAAGATACAAGC | GRMZM2G001696 | PCK | | 3' end |
| JSC4-6 | AGATGGTCATCATGGGCACG | GRMZM2G001696 | PCK | 450 | CDS |
| JSC4-7 | ATGGTATCTTGGCGTTGGGG | GRMZM2G001696 | PCK | | CDS |
| JSC4-8 | AGGGATGGACACTACTTGCG | GRMZM2G085019 | NADP-ME | 790 | CDS |
| JSC4-9 | AATACGTCTGCTCTGCCAGG | GRMZM2G085019 | NADP-ME | | CDS |
| JSC4-10 | AGAACTGCATGTACACTCCCG | GRMZM2G085019 | NADP-ME | 286 | 3' end |
| JSC4-11 | GAGATCTGACTCGTCCAGCC | GRMZM2G085019 | NADP-ME | | 3' end |
